# Oncogenic and Tumor Suppressor Functions for Lymphoid Enhancer Factor 1 in a Murine Model of T Acute Lymphoblastic Leukemia

**DOI:** 10.1101/2021.11.08.467708

**Authors:** Tiffany Carr, Stephanie McGregor, Sheila Dias, Mihalis Verykokakis, Michelle M. LeBeau, Hai-Hui Xue, Elizabeth T. Bartom, Barbara L. Kee

**Author notes:** Address Correspondence to: Barbara L. Kee, Department of Pathology, The University of Chicago, 924 E. 57^th^ Street, JFK Rm 318, Chicago IL, 60637.

## Abstract

T lymphocyte acute lymphoblastic leukemia (T-ALL) is a heterogeneous disease affecting T cells at multiple stages of their development and is characterized by frequent genomic alterations. The transcription factor LEF1 is inactivated through mutation in a subset of T-ALL cases but elevated LEF1 expression and activating mutations have also been identified in this disease. Here we show, in a murine model of T-ALL, that the developmental timing of *Lef1* mutation impacts its ability to function as a cooperative tumor suppressor or oncogene. T cell transformation in the presence of LEF1 allows leukemic cells to become addicted to its presence. In contrast, deletion prior to transformation both accelerates leukemogenesis and results in leukemic cells with altered expression of genes controlling receptor signaling pathways. Our data demonstrate that the developmental timing of *Lef1* mutations impact its apparent oncogenic or tumor suppressive characteristics and demonstrate the utility of mouse models for understanding the cooperation and consequence of mutational order in leukemogenesis.

## Introduction

T acute lymphoblastic leukemia (T-ALL) is an aggressive malignancy that accounts for approximately 10-15% of pediatric and 25% of adult leukemia ^1^. This disease arises in thymocytes and leads to an enlarged thymus causing respiratory distress; however, leukemic cells can disseminate through the blood and infiltrate tissues such as the bone marrow, spleen and liver. Treatment involves intensive chemotherapy but 25% of pediatric cases and nearly 50% of adult cases show therapy resistance or relapse within 5 years ^1^. T-ALL is a heterogeneous disease characterized by transformation of cells at multiple stages of T cell differentiation including ETP-ALL, pro-T, pre-T, cortical and mature ^2,3^. Distinct subsets of T-ALL can be identified by their unique gene expression signatures, genetic alterations, and responses to therapy ^1,4,5^. Recent analysis further highlights that clonal evolution and the progressive mutation contribute to disease evolution ^6^. Understanding the molecular pathogenesis of T-ALL is crucial for the improved development of prognostic markers and tailored therapeutic approaches.

The most broadly occurring mutations in T-ALL affect the Notch signaling pathway. Mutations in the *Notch1* gene occur in > 60% of cases ^7^. Mutations occur within the heterodimerization domain, resulting in ligand independent activation of Notch1, and the intracellular PEST domain, resulting in stabilization of the transcriptionally active form (intracellular Notch1/ICN). In another 15% of T-ALL, mutations occur in *Fbxw7*, which encodes an ubiquitin ligase involved in the PEST-domain dependent degradation of ICN, and other oncogenic proteins such as c-myc ^8-10^. Further, epigenetic alterations affecting the *Notch1* gene can impact the site of transcription initiation and splicing and result in ligand independent activation of Notch1 ^11-13^. Consistent with the oncogenic role of Notch1, ectopic expression of constitutively active forms of Notch1 in mouse T cell progenitors leads to their transformation ^14^.

Activation of the *Tal1* and *Lyl1* genes is also associated with subsets of T-ALL ^4^. Tal1 and Lyl1 are basic helix-loop-helix proteins that bind DNA in association with the E protein transcription factors ^15^, which are critical transcription factors for T cell development ^16^. While Tal1:E protein and Lyl1:E protein dimers have targets implicated in T-ALL ^17-21^, their ability to inhibit E protein homodimer formation is sufficient to promote T cell transformation as revealed by the development of T-ALL like disease in *E2a*^*-/-*^ mice and in mice ectopically expressing inhibitors of E protein DNA binding ^22-25^. *E2a*^*-/-*^ leukemias are characterized by recurrent mutations in the *Notch1* gene, primarily in the PEST domain, as well as altered splicing and transcription initiation leading to ligand independence and these cells require Notch1 for their survival ^11,26^.

A critical target of Notch1 in many cases of T-ALL is c-*myc* ^27-29^. However, in *E2a*^*-/-*^ leukemias Notch1 signaling is essential but it regulates expression of *Lef1*, encoding a TCF1-related transcription factor that is an effector of the Wnt signaling pathway ^26,30,31^. LEF1 appears essential for the survival of *E2a*^*-/-*^ leukemias since siRNA-mediated knockdown of *Lef1* causes cell cycle arrest and the death of leukemias in vitro ^30^. Other murine models of T-ALL, including those arising in *Tcf7*^*-/-*^ and *Ikzf1*^*-/-*^ mice also show increased expression of *Lef1* ^32,33^. Recently, a study of childhood ALL, including 28 patients with T-ALL, revealed a positive prognostic value to high *LEF1* expression and a second study confirmed this for specific LEF1 isoforms ^34,35^. However, in a study of adult T-ALL, 25% of patients had elevated expression of LEF1 that was associated with a poor prognosis ^36^. In 4 patients, two mutations in *LEF1* (K86E and P106L) were found to augment the transcriptional capacity of LEF1. In contrast to these findings of increased LEF1 expression or function, a subset of human T-ALL (18-27%) have inactivating mutations within the *LEF1* gene ^5,37^. These observations suggest that LEF1 can function as both a pro- and anti-leukemia factor but a molecular understanding of the basis for these distinct functions is currently unknown.

To gain further insight into the role of LEF1 in T-ALL we investigated the requirement for LEF1 in the generation of *E2a*^*-/-*^ T cell leukemias. We demonstrate that *E2a*^*-/-*^ leukemias that arose in mice sufficient for LEF1 became dependent on this transcription factor for their survival. However, deletion of *Lef1* prior to T cell transformation resulted in significant alterations in T cell development and a reduced latency to leukemic morbidity. Leukemic cells arising in the latter context resembled *E2a*^*- /-*^ leukemias in that they had mutations in Notch1 and dependence on the Notch signaling pathway as well as frequent trisomy of chromosome 15, and hence increased c-*myc* expression. However, they differed from *E2a*^*-/-*^ leukemias in that they had a CD4^lo^CD8^lo^ phenotype with increased peripheral cell numbers at the time of sacrifice. Cell lines generated from these leukemias reveal differences in expression of multiple genes associated with monocarboxylic transport and hedgehog signaling that could also impact T cell receptor signaling. Our study describes novel models for studying LEF1 function in T-ALL and indicate that LEF1 is a modulator of leukemic transformation providing both addictive and inhibitory functions depending on its availability during transformation.

## Methods

### Mice

All mice used in this study were backcrossed onto an FvB/NJ background for at least 8 generations. All experiments were performed in compliance with the University of Chicago Institutional Animal Care and Use Committee. *E2a*^*-/-*^, *Lef1*^*f/f*^, and Lck-Cre mice and genotyping protocols were described previously ^38^. FvBn/J mice and Lck-Cre mice were purchased from the Jackson Laboratory.

### Flow cytometry

Thymocytes or splenocytes were dissected and dispersed using frosted glass slides followed by filtration through a 100 mM cell strainer. Cells were stained at a concentration of 2 × 10^7^ cells/ml in FACS buffer (PBS + 5% FCS + .02% azide) after incubation with FcBlock. Intracellular staining was performed using the FoxP3/Transcription Factor Staining Kit. The concentration antibody was determined by titration on thymus or spleen. Antibodies were purchased from BioLegend, eBiosciences or Fischer Scientific and specific antibody clones and fluorochromes are available upon request. Antibodies used: Lef1, CD4, CD8, CD117, CD25, CD44, TCRb, CD11b, CD11c, DX5, B220, Gr1. Data was acquired on an LSRII or Fortessa and analyzed using FlowJo (TreeStar).

### Cell Lines

Leukemia cell lines were generated by culturing thymic cells from moribund mice in OPTI-MEM media containing 10% FCS, 2-mercaptoethanol and Pen/Strep/Glu for greater than 2 weeks. All established lines were frozen in 5% DMSO/50%FCS in liquid nitrogen for long term storage.

### In vitro culture

OP9-DL1 Stromal cells were maintained in OPTI-MEM and plated 1 day before used to achieve a near confluent monolayer of cells. Multipotent progenitors were isolated as Ter119-Gr1-cells from e13 fetal liver and cultured on OP9-DL1 in the presence of 5 ng/ml Flt3 ligand, IL-7 and CD117 ligand.

### Retroviral Transduction

The retroviral vectors MigR1, MigR1-Cre, MigR1-DNMAML were described previously ^39^. Retroviral plasmid DNA was isolated using CsCl. Retroviral supernatants were produced by transfecting plasmid DNA into Phoenix cells using Ca_2_PO_4_ precipitation and cells were transduced with retrovirus as previously described ^40^.

### RNA extraction, sequencing and analysis

RNA was extracted from cell lines using Trizol Reagent. RNA was sequenced on a Next-Seq500 and analyzed as described previously^41^. Raw sequence reads were trimmed using Trimmomatic v0.33 (TRAILING:30 MINLEN:20) and then aligned to mouse genome assembly mm10 with TopHat v 2.1.0. Reads were assigned to genes using the htseq-count tool from HTSeq v 0.6.1 and gene annotations from Ensembl release 78. The R package EdgeR was used to normalize the gene counts and to calculate differential expression statistics for each gene for each pairwise comparison of sample groups. Metascape analysis was performed on differential gene expression lists. Genes were considered differentially expressed if the fold change was >2 and false discovery rate <0.05.

### Quantitative PCR

RNA was isolated from sorted or cultured cells using Trizol Reagent (Invitrogen) or the RNAeasy minikit (Qiagen) and was reverse transcribed with Superscript III (Invitrogen). Quantitative RT-PCR was performed in an iCycler (Bio-Rad Laboratories) with SYBR Green (Bio-Rad Laboratories). Expression values were normalized to Hprt and we calculated by the ΔC_T_ method. Primer sequences are available on request.

### Spectral Karyotyping (SKY) Analysis

To characterize the cytogentic pattern of T cell leukemias from *E2a*^*-/-*^*Lef1* ^*Δ/Δ*^ mice, SKY analysis was performed using the ASI SkyPaintTM assay for mouse chromosomes as described previously ^42^ on cell lines or fresh leukemic cells from moribund mice with leukemia (10 metaphase cells were analyzed per case). Karyotype results are in Table S1.

### Western Blot

Total protein extracts were prepared and analyzed by Western blot analysis as described previously ^30^. Primary antibodies used were anti-Notch1 antibody (V1744) reactive with the cleaved cytoplasmic domain (Cell Signaling Technology) and anti-actin (Abcam).

### Data Sharing Statement>

For original data please contact. RNA-sequencing data can be accessed in the Gene Expression Omnibus under GSE186420.

## Results

### LEF1 is required for the survival of E2A-deficient T cell leukemias

Our previous studies revealed that Notch1 is mutated in *E2a*^*-/-*^ T cell leukemias and required for their survival ^26^. We identified *Lef1* as a target of the Notch signaling pathway in these cells and demonstrated that siRNA directed against *Lef1* reduced the viability of these leukemic cells ^30^. Here, using flow cytometry, we found that LEF1 and TCF1 protein can be detected in *E2a*^*-/-*^ leukemias and that LEF1, but not TCF1, was reduced after treatment of cells with a γ-secretase inhibitor, which antagonizes Notch signaling (Fig. 1A, B). To rigorously demonstrate that LEF1 was required for survival of these leukemic cells, we created *E2a*^*-/-*^ mice that were homozygous for alleles of *Lef1* with loxp sites flanking the DNA binding domain ^32^. We generated 2 cell lines from the leukemias arising in these mice and used a retrovirus producing Cre to delete the DNA binding domain of LEF1 (Fig. 1C). The retrovirus produces GFP in addition to Cre and therefore we could track Cre expressing cells by their expression of GFP (Fig. 1C, D). LEF1 protein was lost from a subset of cells after transduction with MigR1-Cre and the frequency of LEF1 negative cells mirrored the frequency of GFP expressing cells (Fig. 1D). QPCR analysis of sorted GFP^+^ cells from MigR1 or MigR1-Cre transduced cells revealed decreased *Lef1* mRNA after transduction of *E2a*^*-/-*^*Lef1*^*f/f*^ but not *E2a*^*-/-*^ leukemias with MigR1-Cre (Fig. 1E). We tracked the fate of cells with deletion of *Lef1* by following the frequency of GFP^+^ cells in the population. GFP^+^ cells in MigR1-Cre expressing *E2a*^*-/-*^ *Lef1*^*f/f*^ cells declined steadily over time consistent with the loss of cells that lacked LEF1 (Fig. 1F). An *E2a*^*-/-*^ leukemia line that was heterozygous for the *Lef1*^*f*^ allele (*E2a*^*-/-*^*Lef1*^*f/+*^) and transduced with MigR1-Cre also showed reduced representation of GFP^+^ cells with time in culture, although not to the degree of *E2A*^*-/-*^*Lef1*^*f/f*^ leukemias (Fig. 1F). In contrast, the same cell lines transduced with MigR1, which does not promote deletion of *Lef1*, demonstrated a stable frequency of GFP^+^ cells over time (Fig. 1D-F). Similarly, *E2a*^*-/-*^ lines transduced with either MigR1 or MigR1-Cre showed stable GFP expression (Fig. 1F). These data demonstrate that LEF1 is required for the maintenance of *E2a*^*-/-*^ T cell leukemia lines in vitro and are consistent with our previous studies using *Lef1* siRNA ^30^.

**Figure 1:**
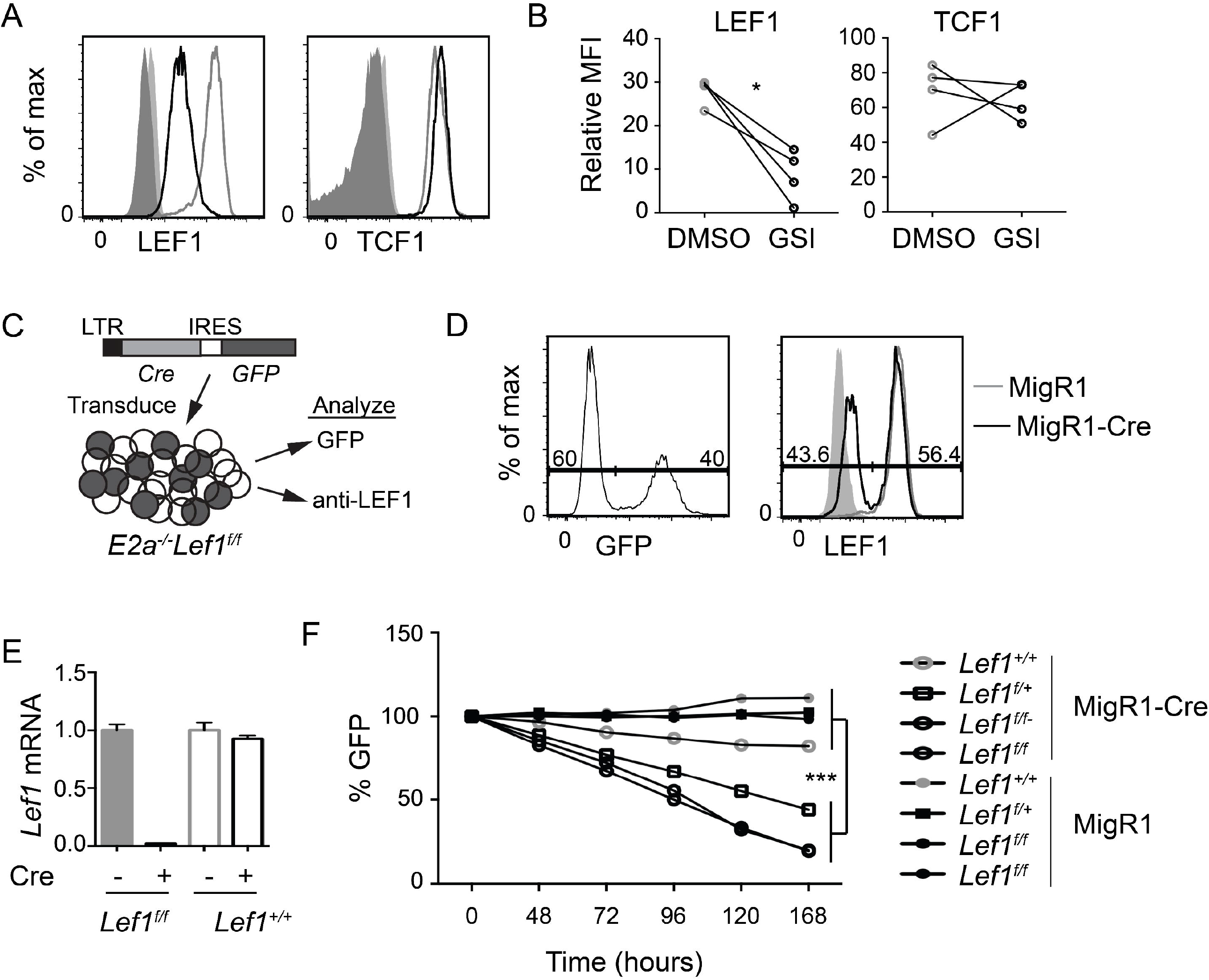
LEF1 is required for the survival of established *E2a*^*-/-*^ T cell leukemias. (A) Flow cytometry for LEF1 (left) or TCF1 (right) in an *E2a*^*-/-*^ T cell leukemia line treated with DMSO (grey) or the Notch1 inhibitor GSI (black). Shaded histograms are isotype control staining. n=4 (B) Summary of the MFI for LEF1 or TCF1 in multiple *E2a*^*-/-*^ leukemia lines. (C) Schematic representation of experiment to inactivate *Lef1* in *E2a*^*-/-*^ *Lef1*^*f/f*^ T cell leukemia. (D) Expression of GFP (left) or Lef1 (right) in *E2a*^*-/-*^*Lef1*^*f/f*^ leukemias transduced with Migr1-Cre (black). For LEF1 staining, Migr1 transduced cells (grey) and isotype controls (shaded histograms) are also shown. (E) qRT-PCR analysis for *Lef1* mRNA in GFP^+^ cells isolated from *E2a*^*-/-*^ leukemias with *Lef1*^*f/f*^ or *Lef1*^*+/+*^ 96 hours after transduction with MigR1 or MigR1-Cre. Error bars represent standard deviation. (F) The relative percent of GFP expressing cells with time in culture after transduction of *E2a*^*-/-*^ leukemias with *Lef1*^*+/+*^, *Lef1*^*f/+*^ or *Lef1*^*f/f*^ genotype. The leukemias were transduced with MigR1 or MigR1-Cre retrovirus. Data are representative of n=4 (A, B), n=2 (D, E) and n=3 (F) experiments. * p<0.05 and ***p<0.005.

### LEF1 is increased in *E2a*^*-/-*^ DN3 thymocytes and promotes their differentiation

To determine whether the increased expression of LEF1 occurs in pre-leukemic mice, we examined the Lineage negative population of the thymus for expression of LEF1. As expected, there were fewer DN3 thymocytes in *E2a*^*-/-*^ mice compared to control with an increased frequency of CD117^lo^ CD25^int^ cells, previously shown to be innate lymphoid cells (Fig. 2A) ^43,44^. The *E2a*^*-/-*^ DN3 cells expressed substantially more LEF1 than control DN3 thymocytes (Fig. 2B). By qRT-PCR we also found increased expression of *Lef1* mRNA in *E2a*^*-/-*^ DN3 thymocytes (Fig. 2C). DN3 thymocytes fail to develop from *E2a*^*-/-*^ multipotent progenitors cultured in vitro on OP9-DL1 ^43,45^, indicating that these cells are compromised with respect to T cell differentiation. To determine whether LEF1 might provide an advantage to these cells, we used a retrovirus to force multipotent progenitor (MPP) cells to ectopically express LEF1. *E2a*^*-/-*^ MPPs transduced with the MigR1 retrovirus, which produce GFP only, failed to generate DN3 cells in vitro, as expected (Fig. 2D, E). In contrast, *E2a*^*-/-*^ MPPs transduced with LEF1 producing retrovirus generated DN3 cells and an increased frequency of DN2 cells. Notably, even control MPPs transduced with LEF1 producing retrovirus showed increased generation of DN2 and DN3 cells (Fig. 2D, E). This propensity to promote differentiation of control and *E2a*^*-/-*^ MPPs was not dependent on the presence of the β-catenin interaction domain (CAT) as a retrovirus producing a mutant form of LEF1 lacking the CAT domain also supported differentiation (Fig. 2D, E). These data lead us to hypothesize that the increased expression of LEF1 in *E2a*^*-/-*^ DN3 thymocytes aids in their differentiation from more immature progenitors and that LEF1’s essential functions are independent of its interaction with β-catenin.

**Figure 2:**
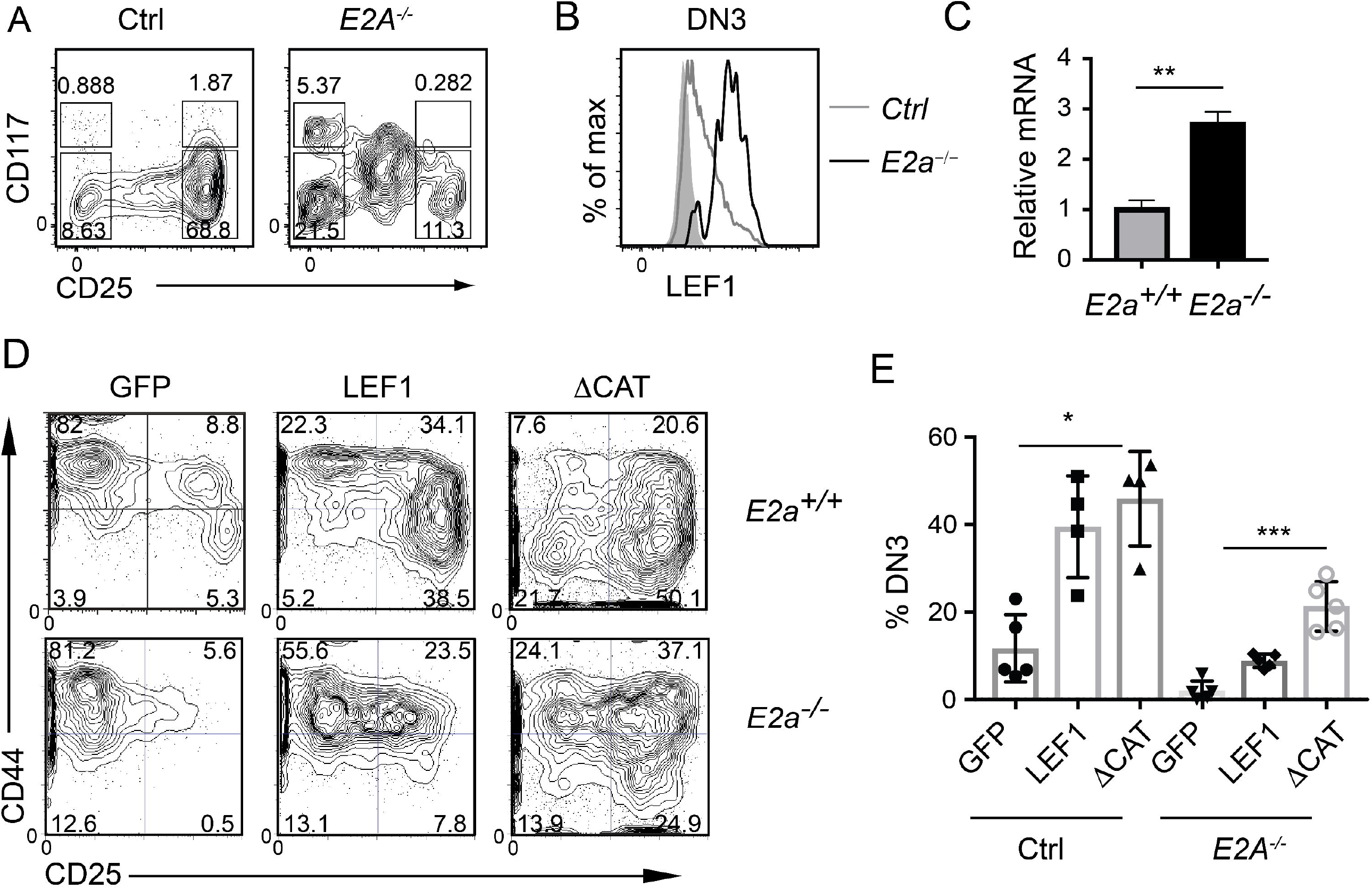
LEF1 is highly expressed in *E2a*^*-/-*^ DN3 cells and promotes their development from MPPs. (A) Flow cytometry showing CD117 and CD25 on Lineage- (CD4, CD8, CD11b, Ter119) thymocytes from Ctrl (left) and *E2a*^*-/-*^ (right) mice. (B) LEF1 expression in DN3 thymocytes from Ctrl (grey) and *E2a*^*-/-*^ (black) mice. (C) Summary of *Lef1* mRNA relative to *Hprt* mRNA in Ctrl (grey) and *E2a*^*-/-*^ (black) DN3 thymocytes. (D) Flow cytometry of MPPs isolated from Ctrl (top) or *E2a*^*-/-*^ (bottom) embryos cultured in vitro for 10 days after retroviral transduction with MigR1 (GFP), MigR1-LEF1 or MigR1-LEF1ΔCAT. Data are gated on GFP^+^ cells. (E) Summary of the %DN3 cells in the Lineage^-^ population of MPPs cultured in vitro as in (D) from the indicated strains. Data are representative of (A, B) n=3, (C) n=2, (D, E) n=4-6 experiments. * p<0.05, **p>0.01 and ***p<0.005.

### T cell specific deletion of *Lef1* in *E2a*^*-/-*^ mice abrogates DN3 development

To test the hypothesis that LEF1 is essential for the development of *E2a*^*-/-*^ DN3 cells and for leukemic transformation, we generated Lck-Cre *E2a*^*-/-*^*Lef1*^*f/f*^ mice, which express Cre starting at the DN2 stage. Lck-Cre^+^ *E2a*^*-/-*^*Lef1*^*f/f*^ mice (DKO) and *E2a*^*-/-*^ mice had similar numbers of total thymocytes and Lineage-negative thymocytes (Fig. 3A-C). However, the Lineage-negative population of DKO mice showed a reduced frequency of DN3 thymocytes (Fig. 3B, D). The total number of DN3 thymocytes was also significantly decreased in DKO compared to *E2a*^*-/-*^ mice (Fig. 3E). In contrast Lck-Cre^+^ *Lef1*^*f/f*^ (*Lef1*^*Δ/Δ*^) mice had thymocyte and Lineage-negative thymocyte numbers, as well as DN3 frequencies that were similar to Control mice (Fig. 3D), indicating that *Lef1* deletion did not have a major impact on these cells.

**Figure 3:**
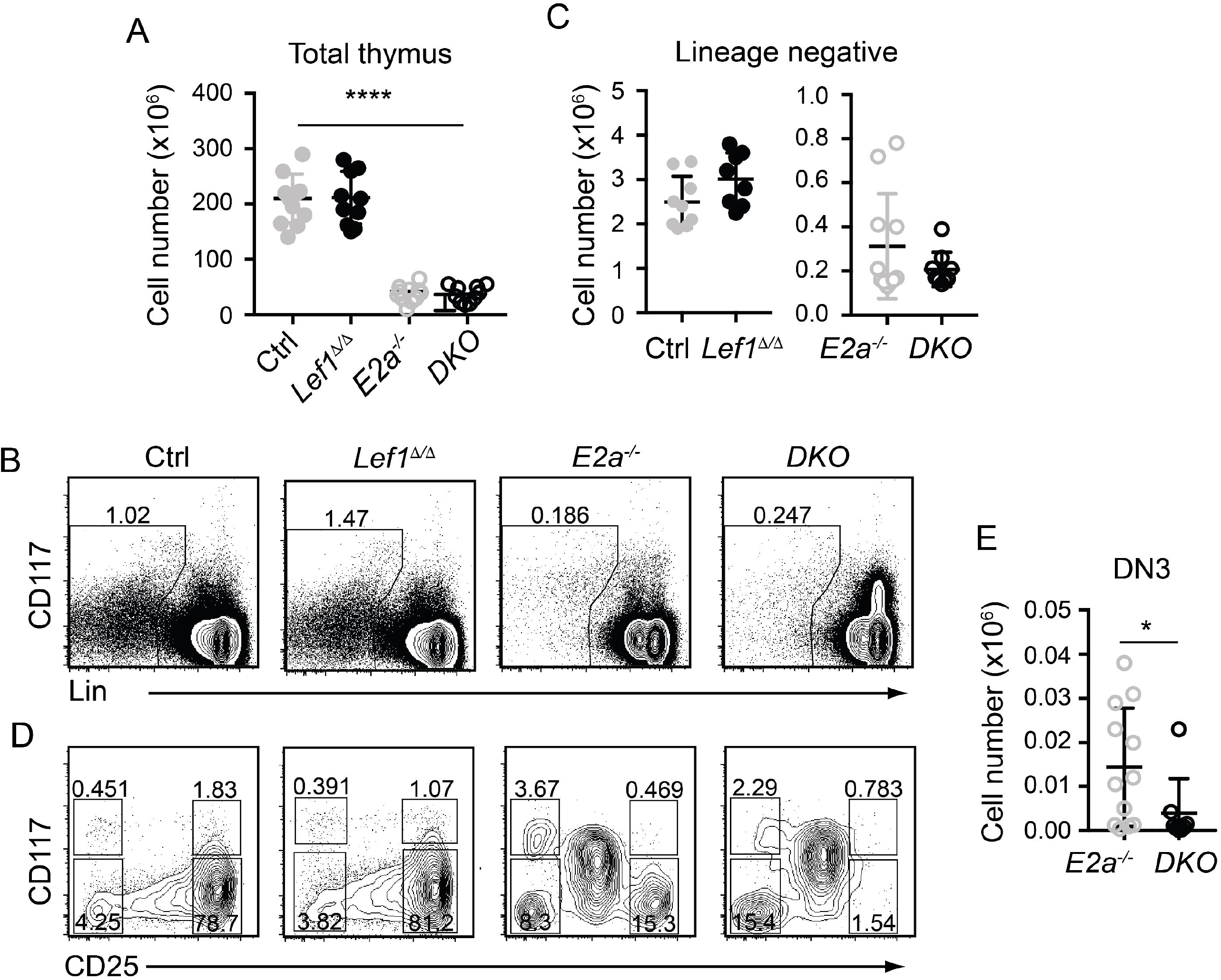
T cell specific deletion of *Lef1* from *E2a*^*-/-*^ mice does not impact thymocyte numbers but results in a loss of DN3 cells. (A) Total thymocyte numbers from mice of the indicated genotype. (B) Flow cytometry showing Lineage markers (CD8, TCRβ, TCRγδ, CD11b, CD11c, NK1.1) and CD117 and the gating strategy for Lin^-^ cells. (C) Lin^-^ thymocyte numbers in mice of the indicated genotypes. (D) Flow cytometry showing CD25 and CD117 on Lin^-^ thymocytes. (E) Summary of DN3 thymocyte numbers in mice of the indicated genotypes. Data is representative on more than 8 experiments *p<0.05. ****p<0.001.

Since the number of thymocytes was similar in *E2a*^*-/-*^ and DKO mice we examined the phenotype of more mature thymocytes. As previously reported, *E2a*^*-/-*^ thymocytes had a reduced frequency of CD4^+^CD8^+^ (DP) thymocytes with and increased frequency of CD4^+^ and CD8^+^ cells. The CD4 by CD8 profile of DKO thymocytes revealed a slightly increased frequency of CD4^+^CD8^+^ (DP) cells and a decreased frequency of CD4^+^ of CD8^+^ single positive thymocytes compared to *E2a*^*-/-*^ mice (Fig. 4A). *E2a*^*-/-*^ mice also have an increased frequency of TCRβ^+^ thymocytes, although the number of these cells is lower than in Ctrl mice (Fig. 4A, B). The frequency and number of TCRβ^+^ cells was reduced in DKO mice compared to *E2a*^*-/-*^ mice, although it was still higher than in Ctrl or *Lef1*^*Δ/Δ*^ mice (Fig. 4A, B). These data suggest that deletion of LEF1 had a subtle but significant impact on the differentiation of thymocytes in the absence of *E2a*. Interestingly, a substantial portion of DKO Lineage^+^ thymocytes, which are primarily DP cells, expressed CD25 (Fig. 4C, D). The frequency of Lineage^+^CD25^+^ cells was also higher in *E2a*^*-/-*^ thymocytes than in Ctrl or *Lef1*^*Δ/Δ*^ thymocytes but was not significantly elevated in number (Fig. 4C, D). Indeed, a direct comparison of DP thymocytes revealed a substantial increase CD25 on DKO compared to *E2a*^*-/-*^ thymocytes (Fig. 4E). Given that CD25 is a known Notch1 target gene ^46^, we tested the hypothesis that Notch1 was expressed in DP thymocytes from DKO cells. By QPCR analysis we observed mRNA for *Notch1* and the Notch1 target gene *Nrarp* in DP thymocytes from DKO but not *E2a*^*-/-*^ or Ctrl mice (Fig. 4F). These data indicate that Notch1 is activated in DKO DP thymocytes.

**Figure 4:**
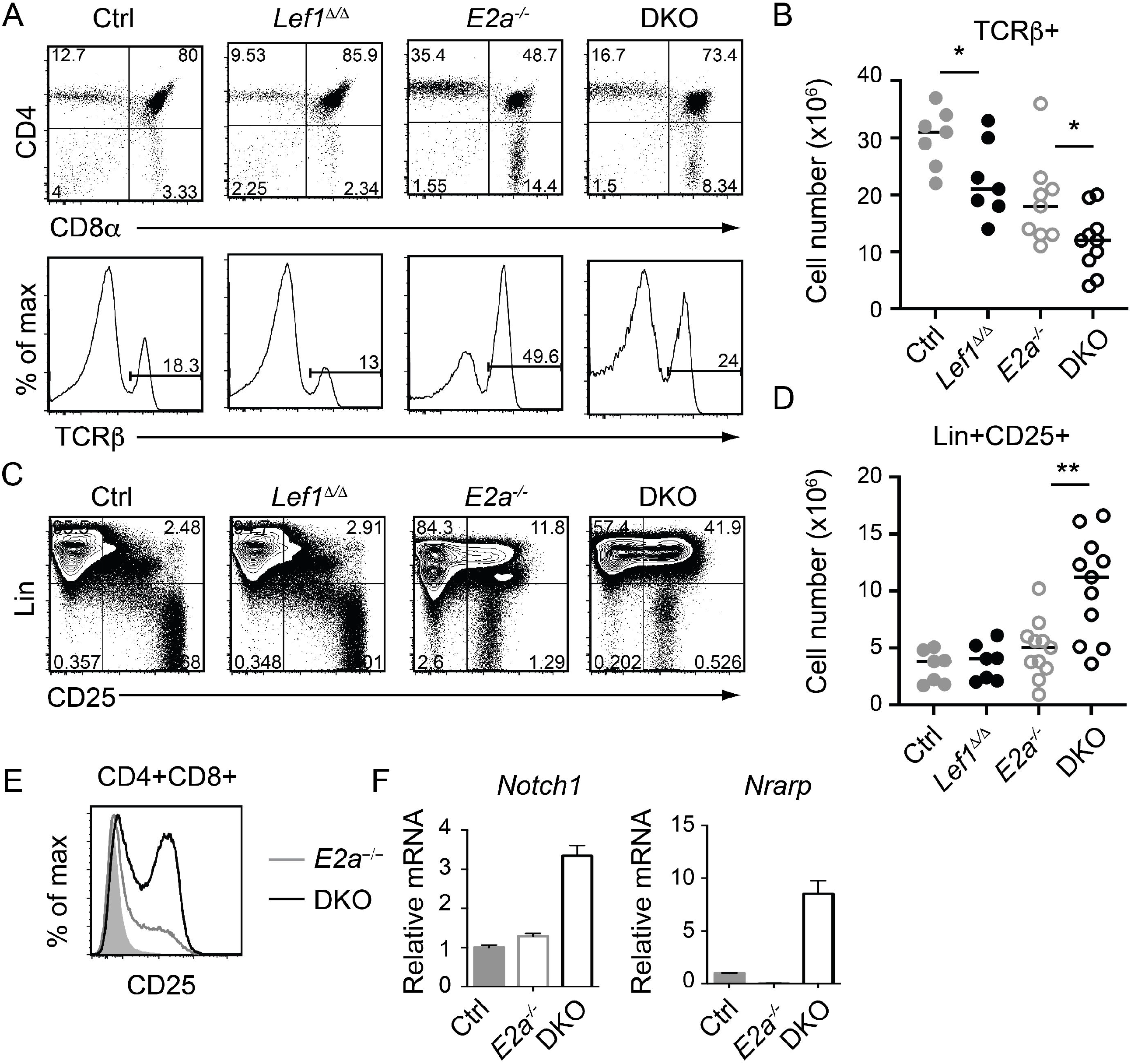
LEF1 restrains expression of CD25, *Notch1*, and *Nrarp* in *E2a*^*-/-*^ CD4^+^CD8^+^ thymocytes. (A) Flow cytometry showing CD4 and CD8 (top) or TCRβ (bottom) on thymocytes from mice of the indicated genotype. (B) Summary of the number of TCRβ^+^ thymocytes in the indicated strains. (C) Flow cytometry showing expression of Lineage markers (CD4, CD8 CD11b, CD11c, CD3e) and CD25. (D) Total number of Lin^+^CD25^+^ cells in mice of the indicated genotype. (E) Flow cytometry of CD25 on CD4^+^CD8^+^ thymocytes from WT (shaded), *E2a*^*-/-*^ (grey) and DKO (black) mice. (F) qRT-PCR for *Notch1* and *Nrarp* on sorted CD4^+^CD8^+^ thymocytes from mice of the indicated genotype. Data is representative of >6 mice (A-D) or 2 experiments (E,F). *p<0.05, **p<0.01.

### DKO mice develop T cell leukemia with reduced latency

To determine whether deletion of *Lef1* impacted the transformation potential of *E2a*^*-/-*^ thymocytes, we allowed the mice to age and monitored them for signs of leukemia. Surprisingly, DKO mice became moribund with an average latency of 100 days (range 80-130 days) whereas the average latency for *E2a*^*-/-*^ mice was 130 days (range 100-170 days), and notably, all mice that we followed developed disease (Fig. 5A). At sacrifice, leukemia was confirmed by counting thymocyte numbers and by flow cytometry of multiple tissues. Cell numbers were similar in the thymus of moribund *E2a*^*-/-*^ or DKO mice, although the range was greater for the DKO cells (Fig. 5B). However, splenic lymphocyte numbers were elevated in the moribund DKO as compared to *E2a*^*-/-*^ mice (Fig. 5C). Primary leukemic cells in DKO mice were frequently low for CD4 and CD8 whereas *E2a*^*-/-*^ leukemias were CD4^hi^CD8^hi^ or contained SP cells (Fig. 5D, E). The DKO leukemias also expressed CD25 without CD44, a phenotype that is distinct from that of the majority of *E2a*^*-/-*^ leukemias (Fig. 5E). Taken together, these data demonstrate that, despite the reduced number of DN3 cells, DKO mice developed T cell leukemia with reduced latency compared to *E2a*^*-/-*^ mice and these leukemias had a distinct surface receptor phenotype.

**Figure 5.**
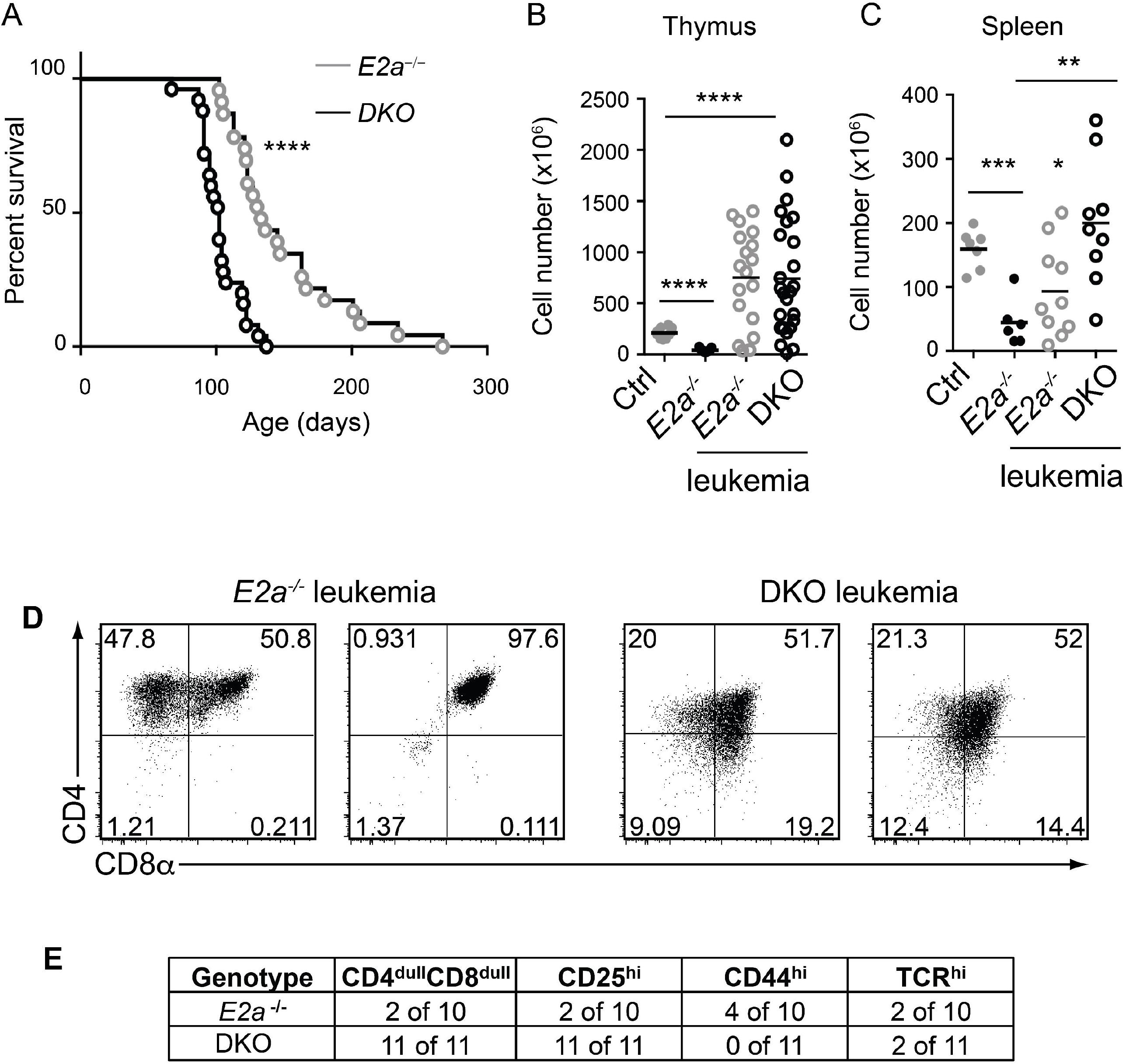
DKO mice have an accelerated onset of T cell leukemia. (A) Kaplan-Meier plot of leukemia incidence in *E2a*^*-/-*^ and DKO mice. (B) Total thymocyte numbers and (C) spleen lymphocyte numbers at time of morbidity in *E2a*^*-/-*^ and DKO mice compared to age matched Ctrl and non-moribund *E2a*^-/-^ mice. (D) Flow cytometry showing CD4 and CD8 on two representative *E2a*^*-/-*^ and DKO thymic leukemias. (E) Summary of phenotype of *E2a*^*-/-*^ and DKO leukemias. **p<0.01, ****p<0.0001

### DKO leukemias have *Notch1* mutations and require Notch signaling

*E2a*^*-/-*^ leukemias have mutations in the *Notch1* gene and they are dependent on Notch signaling for their survival ^26^. To determine whether DKO leukemias also had mutations in the *Notch1* gene, we PCR amplified the 3’ portion of *Notch1* and performed sequencing on 3 DKO cell lines that we established in culture. Notably, we found insertions that resulted in out of frame translation of the PEST domain of Notch1 in all of the DKO lines (Fig. 6A). These mutations are predicted to stabilize ICN1 and, indeed, ICN1 protein could be detected in these leukemias by western blot analysis (Fig. 6B). To determine whether these leukemias were dependent on Notch1 signaling we used retroviral transduction to ectopically express a dominant negative version of the Notch1 co-activator MAML in these cells ^39^. The frequency of cells expressing DN-MAML, identified by their expression of GFP, declined over time in culture regardless of whether they were *E2a*^*-/-*^ or DKO leukemias (Fig. 6C). In contrast, the frequency of GFP^+^ cells remained stable in cultures transduced with the control MigR1 retrovirus (Fig. 6C). Cytogenetic analysis using spectral karyotyping of three DKO primary leukemias revealed trisomy for chromosome 15 containing the *c-Myc* locus, a feature that is also observed in *E2a*^*-/-*^ leukemias (Fig. 6C and Table S1) ^24^. Consistent with this observation, *E2a*^*-/-*^ and DKO cell lines had similar levels of c-*Myc* mRNA (Fig. 6E). These data demonstrate that DKO leukemias, like *E2a*^*-/-*^ leukemias, required Notch signaling for their survival and had mutations impacting the stability of ICN1 expression of c-*Myc*.

**Figure 6:**
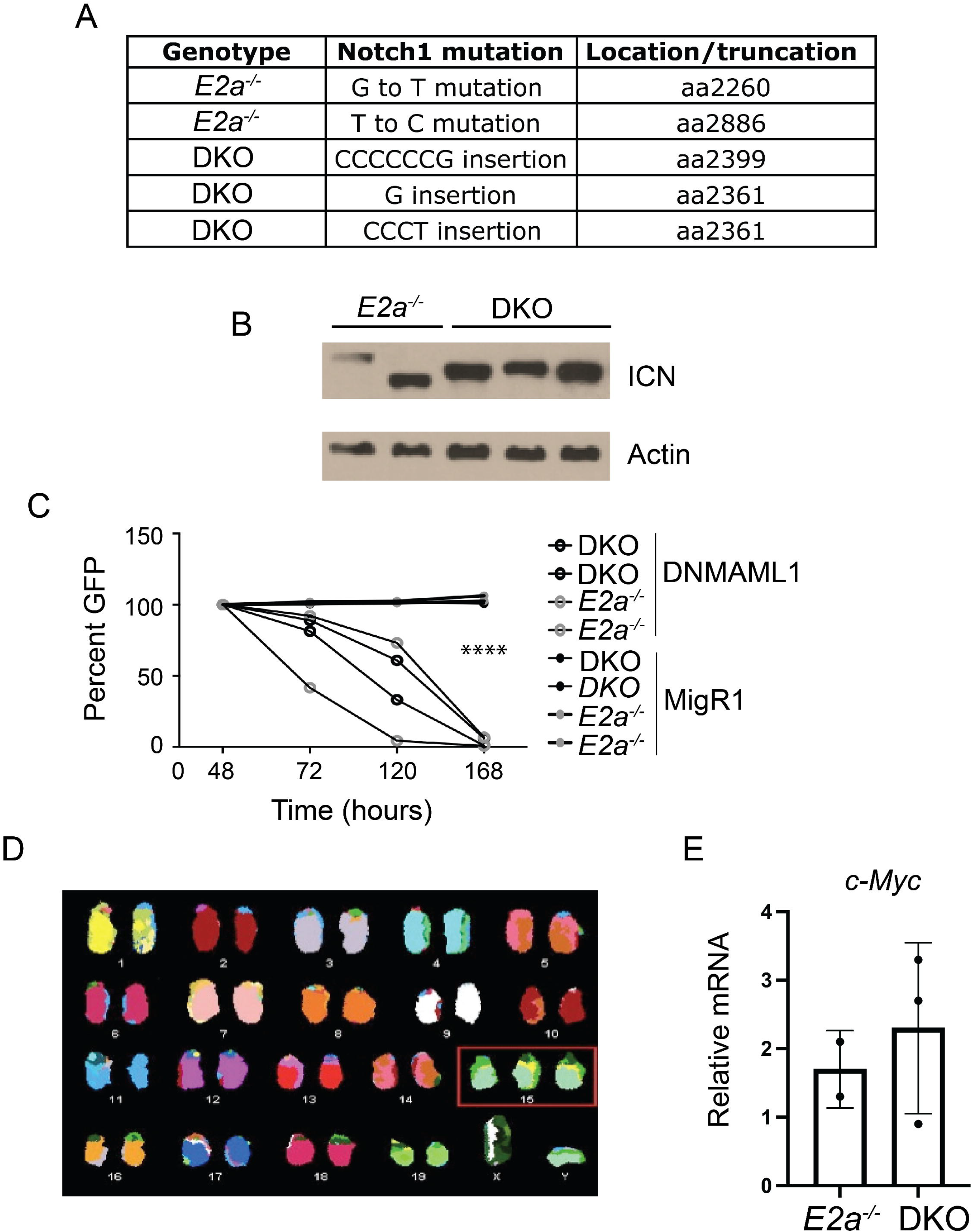
DKO leukemias require Notch1 signaling and have amplification of chromosome 15. (A) Identification of mutations in the *Notch1* gene in two *E2a*^*-/-*^ and 3 DKO leukemias. (B) Western Blot Analysis for cleaved ICN1 in the indicated leukemia lines. Actin is shown as a loading control. (C) Relative GFP expression in leukemias of the indicated genotype after transduction with MigR1 or MigR1-DNMAML. The frequency of GFP^+^ cells was assessed by flow cytometry at the indicated times after transduction. (D) Spectral karyotyping analysis of metaphase cells isolated from a DKO leukemia. (D) qRT-PCR analysis for *c-Myc* in 2 *E2a*^*-/-*^ and 3 DKO leukemias. ****p<0.0001.

### DKO leukemias have an altered transcriptome implicating monocarboxylic acid transport and hedgehog signaling

To gain further insight into the differences between *E2a*^*-/-*^ and DKO leukemias we performed RNA-sequencing on multiple *E2a*^*-/-*^ and DKO leukemia lines. There was a large amount of variation in the gene programs of these leukemias but they were resolved into distinct populations by principle component analysis through PC1 and PC3 (Fig. 7A). We identified 89 genes that were generally decreased in DKO as compared to the *E2a*^*-/-*^ lines and 70 genes that were increased (Log2FC, adj. p-value <0.01) (Fig. 7B). *Tcf7* mRNA appeared to be increased in DKO leukemias but TCF1 protein, unlike LEF1 protein, was expressed similarly in *E2a*^*-/-*^ and DKO when evaluated by flow cytometry (Fig. 7C-and Fig. S1). Analysis of the differentially expressed genes by Metascape revealed that the genes that decreased in the DKO leukemia were enriched for genes involved in IL-4 production whereas those that increased were enriched for genes in the monocarboxylic acid transport (MCT) and the Hedgehog signaling pathways (Fig. 7C, D). Genes in the MCT pathway included *Fabp5, Pla2g12a, Syk* and *Hoxa13* (Fig. 7E). Genes in the Hedgehog signaling pathway included *Axin2, Cdkn1a, Prkch* and *Rasgrp1* (Fig. 7F). Taken together, these data indicate that DKO and *E2a*^*-/-*^ leukemia lines share many common gene expression features but differ in a few key genes that could impact their metabolic requirements or response to external signals.

**Figure 7:**
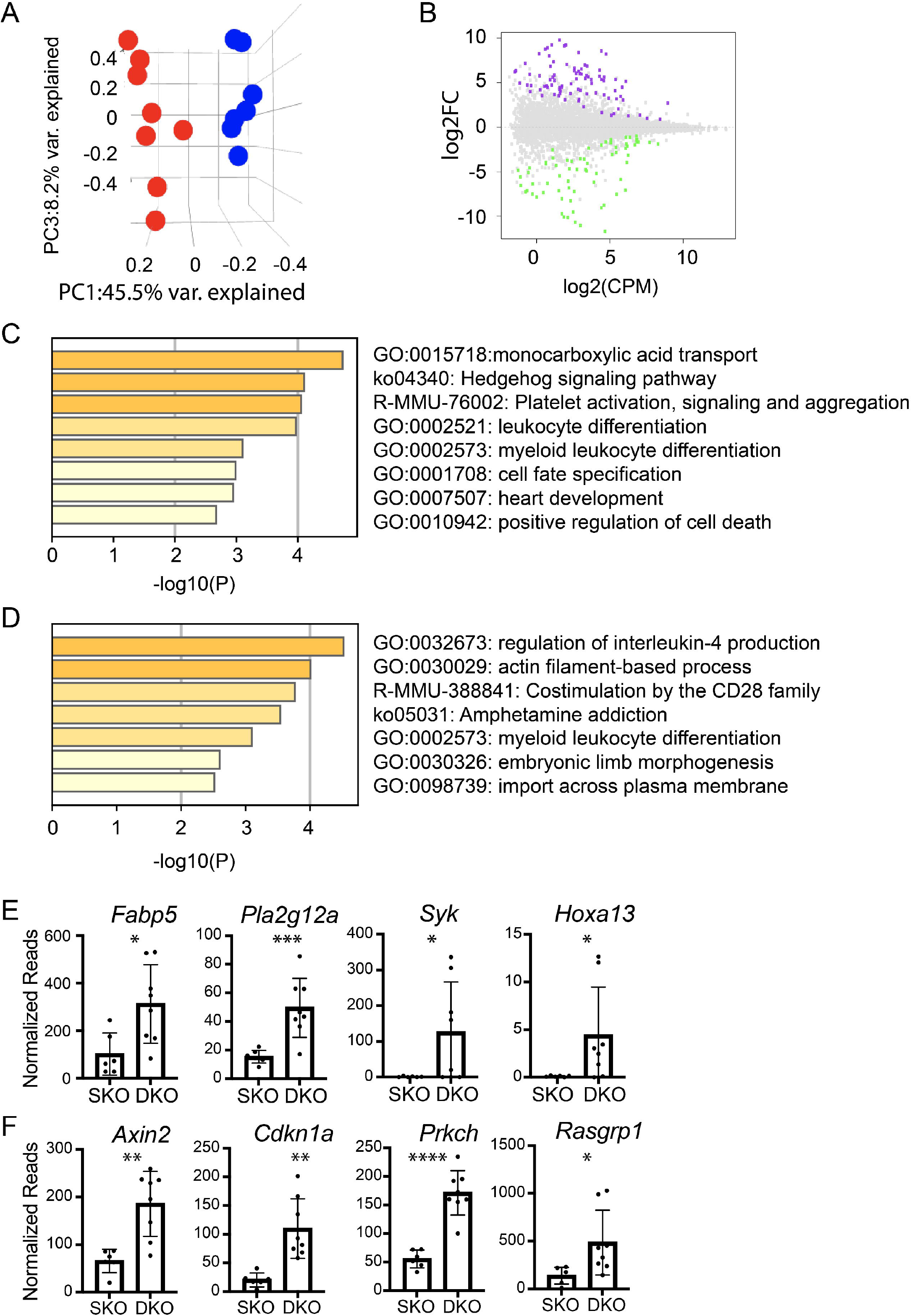
RNA-sequencing of *E2a*^*-/-*^ and DKO leukemia lines reveals differences in multiple genes associated with signaling and monocarboxylic acid transport. (A) Principal Component Analysis of *E2a*^*-/-*^ (blue) and DKO (red) leukemias based on differential gene expression in RNA-seq analysis. (B) MA plot showing Log2FC by Log2(CPM) comparing Rna-seq from E2a-/- and DKO leukemias. Genes that are more highly expressed in *E2a*^*-/-*^ leukemias are indicated in purple and those that are higher in DKO leukemias are indicated in green (adj. p<0.01). (C) Pathway analysis for genes that are increased in *E2a*^*-/-*^ compared to DKO leukemias. (D) Pathway analysis of genes that are increased in DKO compared to *E2a*^*-/-*^ leukemias. (E) Normalized reads for individual *E2a*^*-/-*^ and DKO leukemia samples for genes in the monocarboxylic acid transport pathway or (F) the Hedgehog Signaling Pathway. *adj. p<0.05, **adj. p<0.01, ***adj. p<0.005, and ****adj. p<0.0001.

## Discussion

Over the past 10 years, analysis of gene expression and mutations in T-ALL have revealed both positive and negative associations with *Lef1* ^34,36,37^. Here, we investigated the role of LEF1 in T cell transformation in a murine model of T-ALL. We used conditional alleles of *Lef1* in *E2a*^*-/-*^ T lymphocytes, that were deleted either before or after leukemic transformation and found that LEF1 can function as either an oncogene or a tumor suppressor depending on the context in which it is deleted. When LEF1 is present during the transformation process, leukemias become addicted to its presence. In contrast, when *Lef1* is deleted prior to transformation, early T cell development is altered and there is a more rapid onset of leukemogenesis. We note that leukemias arising in DKO mice had some of the characteristics of human T-ALL with *LEF1* inactivating mutations including mutations in the Notch1 signaling pathway and gain of chromosome 15 ^37^, although these characteristics are shared with *E2a*^*-/-*^ leukemias ^24,26^. However, DKO leukemias had a more a rapid onset than *E2a*^*-/-*^ leukemias mirroring the early development of T-ALL with *Lef1* inactivating mutations ^37^. In addition to providing insight into how LEF1 can be both a positive and negative regulator of leukemogenesis, our data indicate that the order of acquisition of specific mutations in T cell progenitors can impact the phenotype and latency of T-ALL.

The leukemic cells that arise in DKO mice have very dim expression of CD4 and CD8 and express CD25, which distinguishes them from *E2a*^*-/-*^ leukemias. Cell lines established from DKO mice also showed some differences in gene expression from *E2a*^*-/-*^ leukemias. In particular, genes associated with the MCT and Hedgehog signaling pathway, were elevated relative to *E2a*^*-/-*^ leukemias. Among these genes were *Rasgrp1, Pla2g12a* and *Syk*, which are also involved in T cell receptor signaling and their increased expression may be related to the dim expression of CD4 and CD8, which canonically identifies cells undergoing strong TCR signaling or negative selection ^47^. The increased expression of *Fabp5* also raises the possibility that DKO and *E2a*^*-/-*^ leukemias may have different metabolic requirements. While there was quite a bit of variability in the gene programs of individual leukemia lines, these observations raise the possibility that leukemias with different mutation profiles may have unique susceptibilities that could be exploited for therapy.

Lef1 is increased in expression in multiple mouse models of T-ALL. *Tcf7*^*-/-*^ mice also have increased *Lef1* in DN thymocytes but develop an ETP-like T-ALL ^32^. TCF1 was shown to repress *Lef1* and indeed, TCF1 binds to a cluster of sites in the *Lef1* gene and represses transcription from this region in a reporter construct transduced into DN thymocytes. TCF1 is expressed in *E2a*^*-/-*^ thymocytes at levels indistinguishable from control mice indicating that the increased LEF1 is not a consequence of TCF1-deficiency ^43,44^. We showed here that unlike LEF1, TCF1 is not regulated by Notch signaling in *E2a*^*-/-*^ leukemias and therefore we hypothesize that Notch1 regulates *Lef1* directly and independent of TCF1. Deletion of *Lef1* in *Tcf7*^*-/-*^ thymocytes had surprisingly little impact on DN thymocyte development but prevented β-selection and abrogated leukemogenesis ^32^. In contrast, deletion of *Lef1* in *E2a*^*-/-*^ mice did not prevent leukemogenesis. Therefore, TCF1 appears sufficient in *E2a*^*-/-*^ mice to support T cell transformation. TCF1 protein was expressed equivalently in *E2a*^*-/-*^ and DKO leukemias suggesting that LEF1 does not promote leukemogenesis by repressing *Tcf7*. Given that *E2a*^*-/-*^ leukemias are not ETP-like, we do not think that the major function of LEF1 is simply antagonism of TCF1. However, it is possible that high levels of LEF1 impact TCF1 function by competing with TCF1 for binding to TCF1/LEF1 binding sites, either promoting or inhibiting TCF1-like functions. In this scenario, LEF1 may antagonize TCF1 functions that prevent transformation while promoting, or leaving intact, TCF1 functions that support T cell differentiation.

We anticipated that LEF1 would be required for the differentiation of ETPs to the DN3 stage in the absence of E2A and our in vitro experiments supported this hypothesis. Indeed, deletion of *Lef1* in *E2a*^*-/-*^ mice resulted in a loss of DN3 thymocytes. However, total thymocyte numbers were not impacted by deletion of *Lef1* suggesting that LEF1 was not required for T cell development in the absence of *E2a*. Given that we observed increased CD25 and Notch1 signaling in DP thymocytes after deletion of *Lef1*, we hypothesize that LEF1 prevented differentiation of DN3 cells into DP cells. Whether these cells are truly DP thymocytes or DN3 thymocytes that express CD4 and CD8 remains to be investigated. Thus, the altered latency of transformation in DKO mice could be related to differences in the intrinsic susceptibility of thymocytes at different stages of differentiation to transformation in the absence of E2A or to an altered environment in which the transforming progenitors reside.

Taken together, our data demonstrate that LEF1 impacts the developmental trajectory of *E2a*^*-/-*^ T cell progenitors and can act as a tumor suppressor or oncogene depending on its availability during the transformation process. Moreover, our data support the utility of mouse models for understanding the cooperativity and consequence of mutational order on leukemogenesis.

## Supporting information

LEF1 and TCF1 in E2a-/- leukemias

SKY Analysis of leukemia lines

## Acknowledgements

We thank members of the Kee Lab, Warren Pear and Adolfo Ferrando for helpful discussions. Christina Spaulding, Grant van der Voort, Simon Liang, Samantha Cuthbert and Elizabeth M. Davis for technical assistance and Warren Pear for the DN-MAML and Cre producing MigR1 constructs. We are also grateful to the University of Chicago Cytometry and Antibody Technology facility (RRID: SCR_017760) and the Functional Genomics Facility (RRID: SCR_019196) and the Animal Resource Center. This study was supported, in part, by funding from NIAID (R21 AI119894, R21 AI096530 and R01 AI079213) to B.L.K, and (P30 CA014599) to the University of Chicago Comprehensive Cancer Center. RNA-seq data can be found at GSE186420

## Authorship Contributions

T.C., S.M., S.D, and M.V. designed and performed experiments and interpreted data; M.L. oversaw the SKY analysis, H.H.X. provided the *Lef1*^*F/F*^ mice, E.T.B. performed bioinformatics analysis, B.L.K. designed and performed experiments, interpreted data, and wrote the manuscript.

## Conflict-of-interest disclosure

B.L.K. is on the Scientific Advisory Board for Century Therapeutics. All other authors declare no competing interests.

## Figure Legends

**Supplemental Figure 1:** Expression of LEF1 and TCF1 in E2a^*-/-*^ and DKO leukemias. Normalized Reads for *Lef1* (A) and *Tcf7* (B) from RNA-sequencing data using RNA isolated from E2a^-/-^ or DKO leukemia lines. Each dot represents the normalized reads from on line. Flow cytometry for (C) LEF1 and (D) TCF1 in Ctrl thymocytes (left panels) or an E2a^-/-^ (middle panels) or DKO (right panels) leukemia. The shaded histogram is isotype control staining. *** p<0.005.

## Notes

https://www.ncbi.nlm.nih.gov/geo/query/acc.cgi?acc=GSE186420

## References

1. Belver L, Ferrando A. The genetics and mechanisms of T cell acute lymphoblastic leukaemia. Nat Rev Cancer. 2016;16(8):494–507.

2. Bene MC, Castoldi G, Knapp W, et al. Proposals for the immunological classification of acute leukemias. European Group for the Immunological Characterization of Leukemias (EGIL). Leukemia. 1995;9(10):1783–1786.

3. Coustan-Smith E, Mullighan CG, Onciu M, et al. Early T-cell precursor leukaemia: a subtype of very high-risk acute lymphoblastic leukaemia. Lancet Oncol. 2009;10(2):147–156.

4. Ferrando AA, Look AT. Gene expression profiling in T-cell acute lymphoblastic leukemia. Semin Hematol. 2003;40(4):274–280.

5. Noronha EP, Marques LVC, Andrade FG, et al. The Profile of Immunophenotype and Genotype Aberrations in Subsets of Pediatric T-Cell Acute Lymphoblastic Leukemia. Front Oncol. 2019;9:316.

6. Ferrando AA, Lopez-Otin C. Clonal evolution in leukemia. Nat Med. 2017;23(10):1135–1145.

7. Weng AP, Ferrando AA, Lee W, et al. Activating mutations of NOTCH1 in human T cell acute lymphoblastic leukemia. Science. 2004;306(5694):269–271.

8. Malyukova A, Brown S, Papa R, et al. FBXW7 regulates glucocorticoid response in T-cell acute lymphoblastic leukaemia by targeting the glucocorticoid receptor for degradation. Leukemia. 2013;27(5):1053–1062.

9. Thompson BJ, Buonamici S, Sulis ML, et al. The SCFFBW7 ubiquitin ligase complex as a tumor suppressor in T cell leukemia. J Exp Med. 2007;204(8):1825–1835.

10. King B, Trimarchi T, Reavie L, et al. The ubiquitin ligase FBXW7 modulates leukemia-initiating cell activity by regulating MYC stability. Cell. 2013;153(7):1552–1566.

11. Gomez-del Arco P, Kashiwagi M, Jackson AF, et al. Alternative promoter usage at the Notch1 locus supports ligand-independent signaling in T cell development and leukemogenesis. Immunity. 2010;33(5):685–698.

12. Ashworth TD, Pear WS, Chiang MY, et al. Deletion-based mechanisms of Notch1 activation in T-ALL: key roles for RAG recombinase and a conserved internal translational start site in Notch1. Blood. 2010;116(25):5455–5464.

13. Jeannet R, Mastio J, Macias-Garcia A, et al. Oncogenic activation of the Notch1 gene by deletion of its promoter in Ikaros-deficient T-ALL. Blood. 2010;116(25):5443–5454.

14. Allman D, Karnell FG, Punt JA, et al. Separation of Notch1 promoted lineage commitment and expansion/transformation in developing T cells. J Exp Med. 2001;194(1):99–106.

15. de Pooter RF, Kee BL. E proteins and the regulation of early lymphocyte development. Immunol Rev. 2010;238(1):93–109.

16. Anderson MK. At the crossroads: diverse roles of early thymocyte transcriptional regulators. Immunol Rev. 2006;209:191-211.

17. Liau WS, Tan SH, Ngoc PCT, et al. Aberrant activation of the GIMAP enhancer by oncogenic transcription factors in T-cell acute lymphoblastic leukemia. Leukemia. 2017;31(8):1798–1807.

18. Mansour MR, Sanda T, Lawton LN, et al. The TAL1 complex targets the FBXW7 tumor suppressor by activating miR-223 in human T cell acute lymphoblastic leukemia. J Exp Med. 2013;210(8):1545–1557.

19. Sanda T, Lawton LN, Barrasa MI, et al. Core transcriptional regulatory circuit controlled by the TAL1 complex in human T cell acute lymphoblastic leukemia. Cancer Cell. 2012;22(2):209–221.

20. Nagel S, Ehrentraut S, Tomasch J, et al. Transcriptional activation of prostate specific homeobox gene NKX3-1 in subsets of T-cell lymphoblastic leukemia (T-ALL). PLoS One. 2012;7(7):e40747.

21. Kusy S, Gerby B, Goardon N, et al. NKX3.1 is a direct TAL1 target gene that mediates proliferation of TAL1-expressing human T cell acute lymphoblastic leukemia. J Exp Med. 2010;207(10):2141–2156.

22. Kim D, Peng XC, Sun XH. Massive apoptosis of thymocytes in T-cell-deficient Id1 transgenic mice. Mol Cell Biol. 1999;19(12):8240–8253.

23. Morrow MA, Mayer EW, Perez CA, Adlam M, Siu G. Overexpression of the Helix-Loop-Helix protein Id2 blocks T cell development at multiple stages. Mol Immunol. 1999;36(8):491–503.

24. Bain G, Engel I, Robanus Maandag EC, et al. E2A deficiency leads to abnormalities in alphabeta T-cell development and to rapid development of T-cell lymphomas. Mol Cell Biol. 1997;17(8):4782–4791.

25. Yan W, Young AZ, Soares VC, Kelley R, Benezra R, Zhuang Y. High incidence of T-cell tumors in E2A-null mice and E2A/Id1 double-knockout mice. Mol Cell Biol. 1997;17(12):7317–7327.

26. Reschly EJ, Spaulding C, Vilimas T, et al. Notch1 promotes survival of E2A-deficient T cell lymphomas through pre-T cell receptor-dependent and -independent mechanisms. Blood. 2006;107(10):4115–4121.

27. Weng AP, Millholland JM, Yashiro-Ohtani Y, et al. c-Myc is an important direct target of Notch1 in T-cell acute lymphoblastic leukemia/lymphoma. Genes Dev. 2006;20(15):2096–2109.

28. Sharma VM, Calvo JA, Draheim KM, et al. Notch1 contributes to mouse T-cell leukemia by directly inducing the expression of c-myc. Mol Cell Biol. 2006;26(21):8022–8031.

29. Palomero T, Lim WK, Odom DT, et al. NOTCH1 directly regulates c-MYC and activates a feed-forward-loop transcriptional network promoting leukemic cell growth. Proc Natl Acad Sci U S A. 2006;103(48):18261–18266.

30. Spaulding C, Reschly EJ, Zagort DE, et al. Notch1 co-opts lymphoid enhancer factor 1 for survival of murine T-cell lymphomas. Blood. 2007;110(7):2650–2658.

31. Steinke FC, Xue HH. From inception to output, Tcf1 and Lef1 safeguard development of T cells and innate immune cells. Immunol Res. 2014;59(1-3):45-55.

32. Yu S, Zhou X, Steinke FC, et al. The TCF-1 and LEF-1 transcription factors have cooperative and opposing roles in T cell development and malignancy. Immunity. 2012;37(5):813–826.

33. Geimer Le Lay AS, Oravecz A, Mastio J, et al. The tumor suppressor Ikaros shapes the repertoire of notch target genes in T cells. Sci Signal. 2014;7(317):ra28.

34. Jia M, Zhao HZ, Shen HP, et al. Overexpression of lymphoid enhancer-binding factor-1 (LEF1) is a novel favorable prognostic factor in childhood acute lymphoblastic leukemia. Int J Lab Hematol. 2015;37(5):631–640.

35. Erbilgin Y, Hatirnaz Ng O, Can I, et al. Prognostic evidence of LEF1 isoforms in childhood acute lymphoblastic leukemia. Int J Lab Hematol. 2021.

36. Guo X, Zhang R, Liu J, et al. Characterization of LEF1 High Expression and Novel Mutations in Adult Acute Lymphoblastic Leukemia. PLoS One. 2015;10(5):e0125429.

37. Gutierrez A, Sanda T, Ma W, et al. Inactivation of LEF1 in T-cell acute lymphoblastic leukemia. Blood. 2010;115(14):2845–2851.

38. Carr T, Krishnamoorthy V, Yu S, Xue HH, Kee BL, Verykokakis M. The transcription factor lymphoid enhancer factor 1 controls invariant natural killer T cell expansion and Th2-type effector differentiation. J Exp Med. 2015;212(5):793–807.

39. Maillard I, Weng AP, Carpenter AC, et al. Mastermind critically regulates Notch-mediated lymphoid cell fate decisions. Blood. 2004;104(6):1696–1702.

40. Kee BL. Id3 induces growth arrest and caspase-2-dependent apoptosis in B lymphocyte progenitors. J Immunol. 2005;175(7):4518–4527.

41. Jacobsen JA, Woodard J, Mandal M, et al. EZH2 Regulates the Developmental Timing of Effectors of the Pre-Antigen Receptor Checkpoints. J Immunol. 2017;198(12):4682–4691.

42. Le Beau MM, Espinosa 3rd R, Davis EM, Eisenbart JD, Larson RA, Green ED. Cytogenetic and molecular delineaton of a region of chroosome 7 commonly deleted in malignant myeloid diseases. Blood. 1996;88(6):1930–1935.

43. Xu W, Carr T, Ramirez K, McGregor S, Sigvardsson M, Kee BL. E2A transcription factors limit expression of Gata3 to facilitate T lymphocyte lineage commitment. Blood. 2013;121(9):1534–1542.

44. Miyazaki M, Miyazaki K, Chen K, et al. The E-Id Protein Axis Specifies Adaptive Lymphoid Cell Identity and Suppresses Thymic Innate Lymphoid Cell Development. Immunity. 2017;46(5):818–834 e814.

45. Ikawa T, Kawamoto H, Goldrath AW, Murre C. E proteins and Notch signaling cooperate to promote T cell lineage specification and commitment. J Exp Med. 2006;203(5):1329–1342.

46. Gothert JR, Brake RL, Smeets M, Duhrsen U, Begley CG, Izon DJ. NOTCH1 pathway activation is an early hallmark of SCL T leukemogenesis. Blood. 2007;110(10):3753–3762.

47. Gascoigne NR, Palmer E. Signaling in thymic selection. Curr Opin Immunol. 2011;23(2):207–212.

