## Supplementary material for "Oncogenic and Tumor Suppressor Functions for Lymphoid Enhancer Factor 1 in a Murine Model of T Acute Lymphoblastic Leukemia": LEF1 and TCF1 in E2a-/- leukemias

**Supplemental Figure 1:**

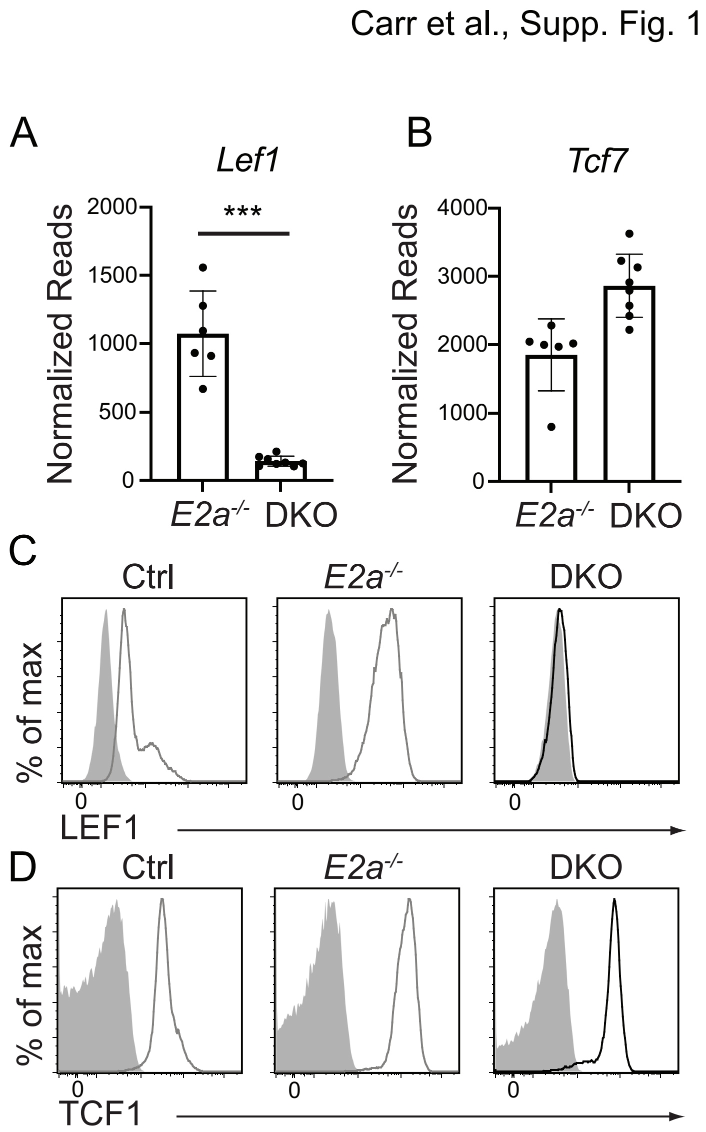


**Supplemental Figure 1:** Expression of LEF1 and TCF1 in *E2a*$^{-/-}$ and DKO leukemias. Normalized Reads for *Lef1* (A) and *Tcf7* (B) from RNA-sequencing data using RNA isolated from E2a$^{-/-}$ or DKO leukemia lines. Each dot represents the normalized reads from on line. Flow cytometry for (C) LEF1 and (D) TCF1 in Ctrl thymocytes (left panels) or an E2a$^{-/-}$ (middle panels) or DKO (right panels) leukemia. The shaded histogram is isotype control staining. *** $p<0.005$.
